# Clemizole and Trazodone are Effective Antiseizure Treatments in a Zebrafish Model of STXBP1 Disorder

**DOI:** 10.1101/2022.01.30.478390

**Authors:** Maia Moog, Scott C. Baraban

## Abstract

**Objective:** CRISPR-Cas9-generated zebrafish carrying a 12 base-pair deletion in *stxbpb1b*, a paralog sharing 79% amino acid sequence identity with human, exhibit spontaneous electrographic seizures during larval stages of development. Zebrafish *stxbp1b* mutants provide an efficient preclinical platform to test antiseizure therapeutics. The present study was designed to test prototype antiepileptic drugs approved for clinical use and two recently identified repurposed drugs with antiseizure activity.

**Methods:** Larval homozygous *stxbp1b* zebrafish (4 days post-fertilization) were agarose-embedded and monitored for electrographic seizure activity using a local field recording electrode placed in midbrain. Frequency of ictal-like events was evaluated at baseline and following 45 min of continuous drug exposure (1 mM, bath application). Analysis was performed on coded files by an experimenter blinded to drug treatment and genotype.

**Results:** Phenytoin, valproate, ethosuximide, levetiracetam, and diazepam had no effect on ictal-like event frequency in *stxbp1b* mutant zebrafish. Clemizole and trazodone decreased ictal-like event frequency in *stxbp1b* mutant zebrafish by 80% and 83%, respectively. These results suggest that repurposed drugs with serotonin receptor binding affinities could be effective antiseizure treatments.

**Significance:** Clemizole and trazodone were identified in a larval zebrafish model for Dravet syndrome. Based primarily on these preclinical zebrafish studies, compassionate-use and double-blind clinical trials with both drugs have progressed. The present study extends this approach to a preclinical zebrafish model representing STXBP1-related disorders, and suggests that future clinical studies may be warranted.

## 1 Introduction

*De novo* mutations in neuronal protein STXBP1 (syntaxin-binding protein 1) represent one of the most common causes of neurodevelopmental disorders and epilepsy. *STXBP1* mutations, estimated at 250 or more, have been described in patients diagnosed with Ohtahara syndrome, Dravet syndrome, Lennox–Gastaut syndrome, West syndrome, and atypical Rett syndrome (1–4). Speech problems, intellectual disability, movement disorders and electroencephalographic (EEG) abnormalities are common clinical presentations associated with these mutations. Seizures in individuals with STXBP1-related disorders, often seen as infantile spasms in the first year of life, evolve to include focal, tonic-clonic and absence seizures over the next several years (5–7). Antiepileptic drug (AED) therapy shows only modest seizure control (largely restricted to the first two years and certain seizure types), but long-term options are generally poor as severe morbidity and high mortality outcomes emerge (8).

These neurodevelopmental disorders are linked to heterozygous missense mutations, nonsense mutations, frame shifts, and deletions in the *STXBP1* gene (6). Haploinsufficient mutations affecting *STXBP1* result in a nonfunctional protein unable to bind synaptobrevin/VAMP2 and synaptosomal-associated protein 25 (SNAP25) e.g., highly conserved neuronal soluble NSF attachment protein receptor complexes (or SNAREs) that drive synaptic transmission via synaptic vesicle exocytosis and secretion of neuropeptides (9, 10). Although SNARE machinery is required for both excitatory (glutamatergic) and inhibitory (GABAergic) synapses, inhibitory synaptic transmission showed strong rundown in cultured hippocampal neurons from *STXBP1*^+/-^ mice (11) and impaired synaptic inhibition mediated by parvalbumin-positive interneurons was reported in cortical slices from *STXBP1*^+/-^ mice (12). These mice also exhibit impairments in cognitive function assessed using a novel object recognition test, hyperactivity in an open-field test and anxiety-like behaviors in elevated plus maze and light-dark chamber tests. Chronic video-EEG recording revealed clusters of spike-wave discharges suggestive of an epileptic phenotype in these animals. STXBP1 mutations modeled in zebrafish include (i) a homozygous *stxbp1b* mutant zebrafish generated by CRISPR/Cas9 editing characterized by spontaneous electrographic seizures (13) and enhanced network cascade activity during interictal periods (14) and (ii) a transgenic mutant overexpressing human STXBP1 (W288X) with spontaneous seizure-like behaviors and increased c-Fos expression (15). Genetically modified zebrafish epilepsy models are effective in identification of novel antiepileptic drugs that have clinical applications (16–20) and the focus of these studies.

## 2 Methods

### 2.1 Zebrafish

Adult male and female zebrafish (TL background strain) were maintained on a 14:10 hour light/dark cycle following standard methods at 28°C. Fish system water conditions were maintained in the following ranges by automated feedback controls: 29-30°C, pH 7.5-8.0, conductivity (EC) 690-710. Zebrafish were housed in rectangular 1.8-L polycarbonate tanks at a density between 4 and 7 adults per tank. Larvae were raised in embryo media consisting of 0.03% Instant Ocean (Aquarium Systems, Inc.) and 0.0002% methylene blue in reverse osmosis-distilled water. Heterozygous *stxbp1b* zebrafish generated by clustered regularly interspaced short palindromic repeats (13) were incrossed and raised in polystyrene petri dishes (100 x 20 mm; FisherBrand #FB0875711Z) to 4 days post-fertilization (dpf). Zebrafish sex cannot be determined until approximately three weeks post-fertilization (21). All procedures followed National Institute of Health and the University of California, San Francisco guidelines and were approved by the Institutional Animal Care and Use Committee (protocol #AN186964-02A).

### 2.2 Electrophysiology

At 4 dpf, zebrafish larvae with darker pigmentation (Fig. 1B) were randomly selected, briefly exposed to cold anesthesia, and immobilized, dorsal side up, in 2% low-melting point agarose (Fisher Scientific) within a slice perfusion chamber (Siskiyou Corporation, Model PC-V). Slice chambers containing two larvae were placed on the stage of an upright microscope (Olympus BX-51W) and monitored continuously using a Axiocam digital camera (Zeiss). Under visual guidance, gap-free local field potential recordings (LFP; 60 min duration) were obtained from optic tectum using a single glass microelectrode (WPI glass #TW150 F-3) filled with 2 mM NaCl internal solution (~1 μm tip diameter) and positioned using a micromanipulator (Siskiyou Corporation, Model MX1641). An Ag/AgCl pellet electrode (Warner Instruments, Part #E205) was used as a submerged bath ground electrode. LFP voltage signals recorded on an Axon MultiClamp 700A amplifier (Molecular Devices) were amplified at a gain of 20x and filtered at 1 kHz (−3 dB; eight-pole Bessel; Cygnus Technology, Inc.), digitized at 10 kHz using a Digidata 1320 A/D interface (Molecular Devices) and stored on a Dell PC computer running AxoScope 10.3 software (Molecular Devices). LFP recordings were initiated ~10 min after agarose embedding. All embedded larvae were continuously monitored for blood flow and heart rate using an Axiocam digital camera via a 4x objective on an Olympus BX-51 upright microscope. Electrophysiology files were coded for *post hoc* off-line analysis.

**Figure 1.**
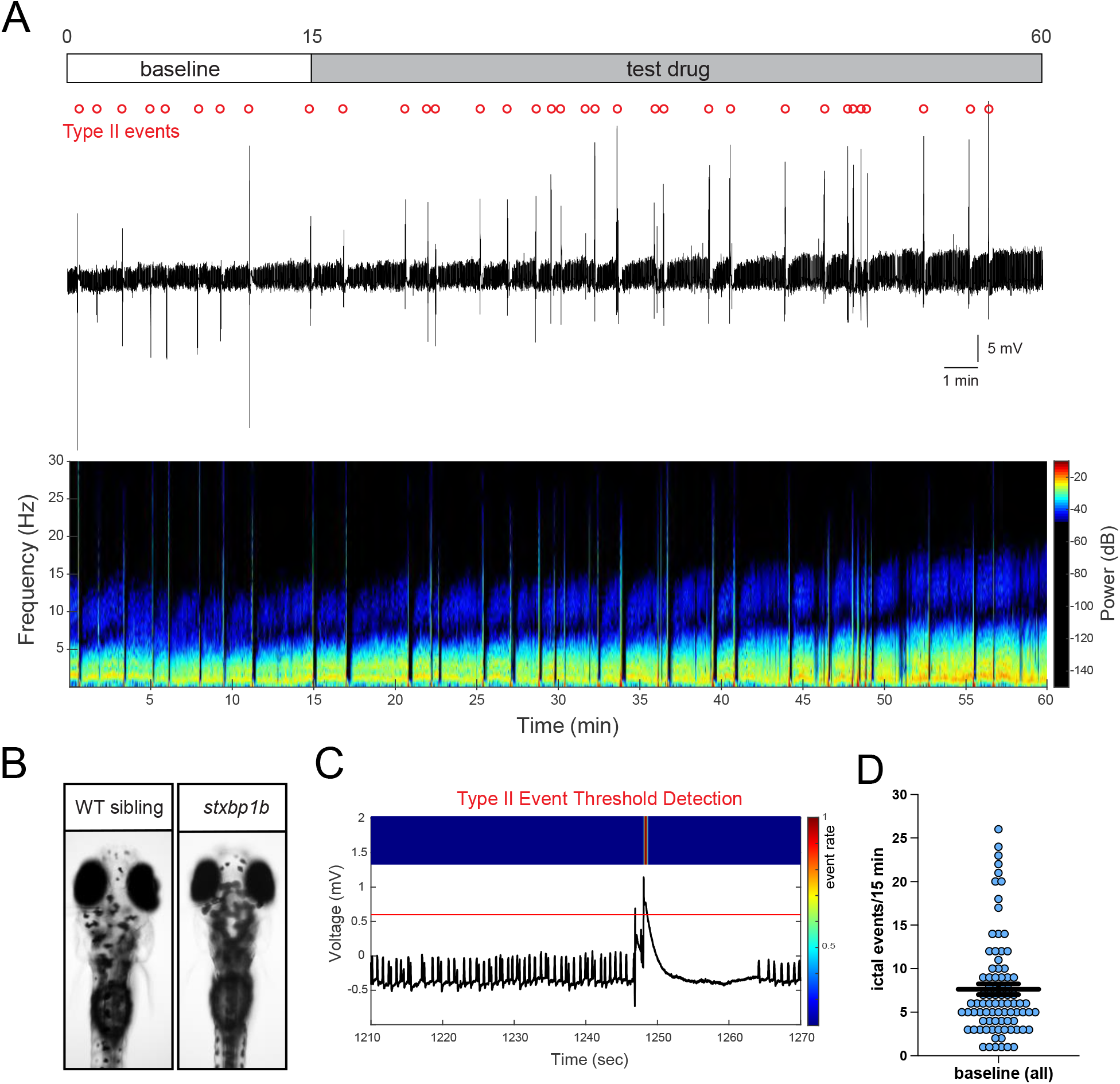
Seizure phenotype in larval *stxbp1b* mutant zebrafish. A, Local field potential recording from a representative agarose-embedded stxbp1b mutant at 4 dpf. Electrode is located in the optic tectum. A 60-min continuous gap-free recording trace (top) and associated spectrogram (bottom) are shown. Type II ictal-like events are denoted by open circles (red). Protocol is shown for baseline (0-15 min) and test drug exposure (15-60 min) periods. B, Representative images from age-matched WT sibling and darkly pigmented stxbp1b mutant larvae at 4 dpf. C, Example of a Type II ictal-like event identified using an amplitude threshold detector set to exclude small-amplitude brief interictal-like activity. Note the presence of a brief period of postictal depression in the electrical recording following the event. D, Plot showing the frequency of Type II ictal-like events in *stxbp1b* mutant larvae used for pharmacology studies (n = 84).

### 2.3 Pharmacology

Drugs were commercially sourced from Millipore-Sigma [phenytoin sodium (PHT), valproic acid (VPA), ethosuximide (ESX), levetiracetam (LEV), diazepam (DZP) and trazodone hydrochloride (TRZ)] or Laurus Labs [clemizole hydrochloride (CLM)]. Drugs chosen for testing were selected from the cohort of U.S. Food and Drug Administration approved clinical treatments for epilepsy. Stock solutions (10 or 50 mM) were made in dimethyl sulfoxide, frozen and freshly diluted in embryo medium for electrophysiology assays. Drug solutions (~0.5 ml) were bath applied at room temperature. All drugs were tested at a concentration of 1 mM as previous studies empirically indicate mM drug concentrations are appropriate for acute pharmacology studies in agarose-embedded larvae (16, 22–28). Experiments were performed on at least three independent clutches of larvae for each drug.

### 2.4 Data analysis

LFP analysis focused on long-duration (> 1 sec), large-amplitude (> 0.5 mV) Type II ictal-like multi- or poly-spike events as these were previously shown to correlate with convulsive seizure behaviors (29) and brain-wide network hypersynchronization (30). Baseline noise level was measured as 0.017 ± 0.005 mV (n = 75). Quantification of Type II event frequency was performed using threshold detection (see Fig. 1C). The number of Type II ictal events was calculated for each larva at baseline and following 45 min of continuous drug exposure. Drugs that suppress Type II ictal events represent potential antiseizure agents (16, 20, 22, 27). All files were un-coded and combined with *post hoc* genotyping data at the end of this process. Data in this manuscript are presented as mean ± standard error of the mean. Student’s t-tests were used for comparisons between baseline and drug treatment groups. Statistical analyses were performed using GraphPad Prism 9 software. A *p* value < 0.05 was considered statistically significant.

## 3 Results

### 3.1 Ictal-like epileptiform activity in *stxbp1b* mutants

As described (13, 14, 29), a zebrafish model of STXBP1-related disorders was generated using CRISPR-Cas9 gene editing. LFP recordings in homozygous *stxbp1b* mutant larvae (4 dpf) confirmed the presence of spontaneous ictal-like (Type II) epileptiform events (Figs. 1A-B). Large-amplitude Type II events were characterized by a brief period of post-ictal depression (Fig. 1C) and a baseline frequency of approximately 0.01 Hz (7.6 ± 0.6 events/15 min; Fig. 1D).

### 3.2 Antiepileptic drugs

STXBP1-related disorders are classified as a drug resistant epilepsy e.g., failure to achieve seizure control with at least two AEDs (31). Using an electrophysiology LFP-based assay, we tested five prototype AEDs (PHT, VPA, ESX, LEV and DZP) covering a wide mechanism of action spectrum (32, 33). Continuous LFP recordings (60 min) were obtained from agarose-immobilized *stxbp1b* mutant larvae. AEDs were bath applied at a concentration of 1 mM. Type II event frequencies were measured at baseline (0-15 min) and following 45 min of drug exposure (45-60 min). No statistically significant changes in event frequency were noted for any of the five AEDs tested (Fig. 2).

**Figure 2.**
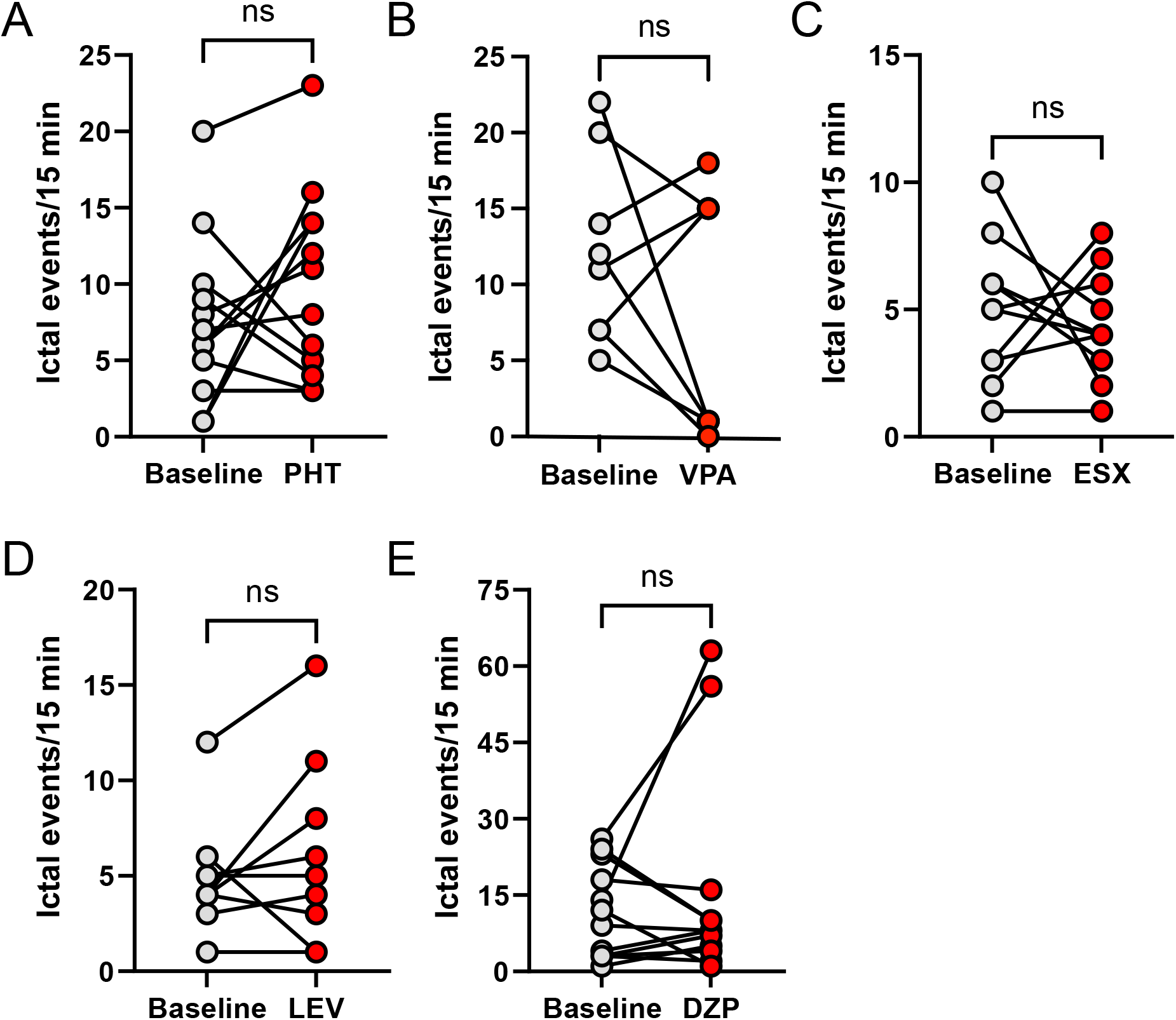
Effect of AEDs on ictal-like seizure events in *stxbp1b* mutant zebrafish. Individual value plots of ictal event frequency at baseline (0-15 min) and 45-60 min after continuous bath application of a test drug. Independent experiment pairs are indicated by the connected line. A, Phenytoin (PHT, n = 13); not significant (ns) *p* = 0.3364. B, Valproate (VPA, n = 8); not significant (ns) *p* = 0.1274. D, Ethosuximide (ESX, n = 11); not significant (ns) *p* = 0.5364. E, Levitracetam (LEV, n = 9); not significant (ns) *p* = 0.5337. E, Diazepam (PHT, n = 12); not significant (ns) *p* = 0.5330. Student’s unpaired *t* tests.

### 3.3 Clemizole and trazodone

Clemizole and trazodone were discovered as repurposed drugs effective in reducing electrographic seizure activity in a Dravet syndrome model e.g., *scn1lab* mutant larvae (16, 17, 20, 34). Continuous LFP recordings (60 min) were obtained from agarose-immobilized *stxbp1b* mutant larvae. Clemizole and trazodone were bath applied at a concentration of 1 mM. Type II event frequencies were measured at baseline (0-15 min) and following 45 min of drug exposure (45-60 min). Event frequencies were significantly reduced following exposure to clemizole (Figs. 3A-B) and trazodone (Fig. 3C-D), by 80% and 83% respectively.

**Figure 3.**
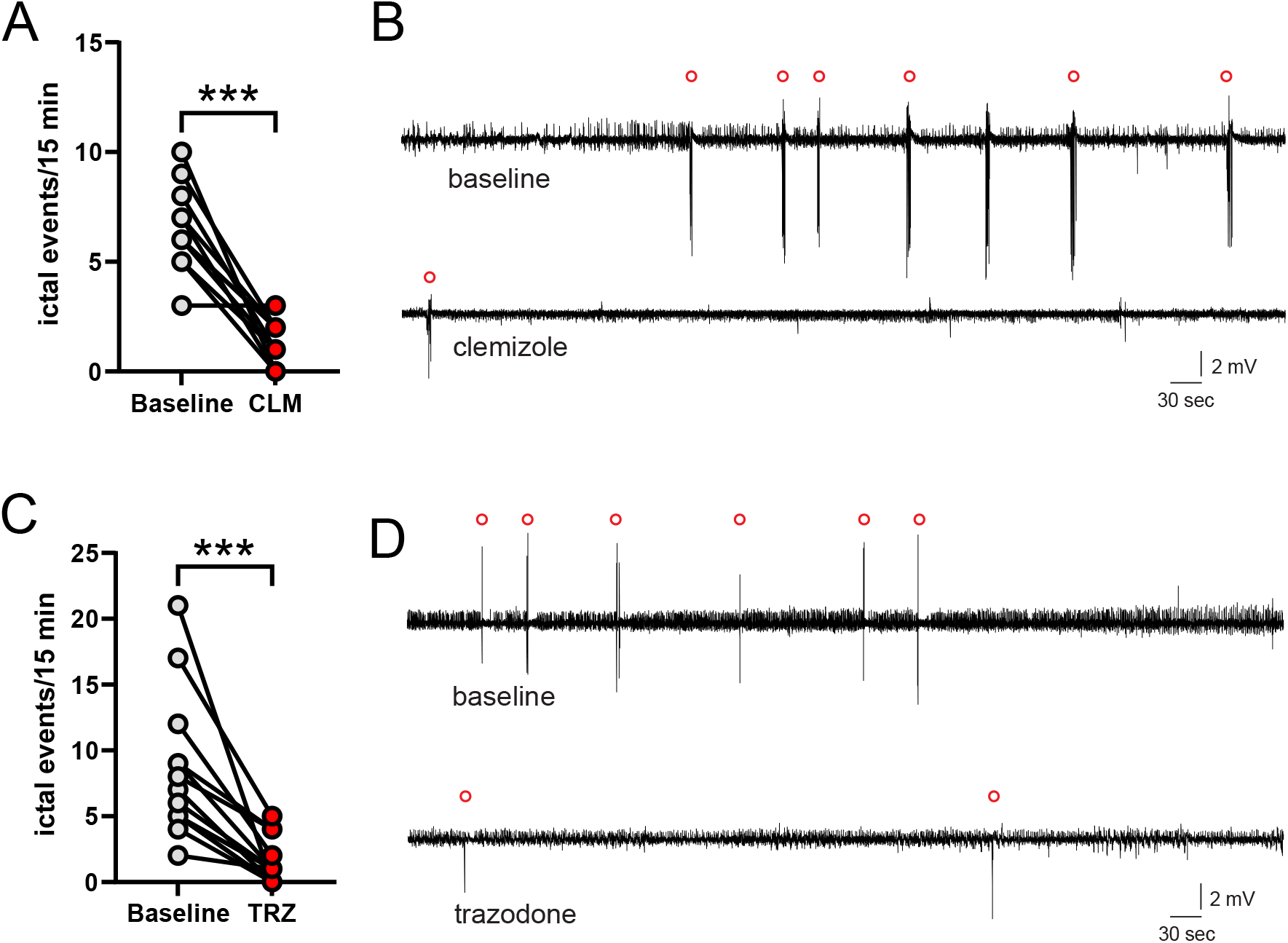
Effect of clemizole and trazodone on ictal-like seizure events in *stxbp1b* mutant zebrafish. A, Individual value plot of ictal event frequency at baseline (0-15 min) and 45-60 min after continuous bath application of clemizole (CLM, n = 11). Independent experiment pairs are indicated by the connected line. \*\*\**p* < 0.0001. B, Representative 60 min LFP traces for *stxbp1b* mutant zebrafish at baseline (6.5 ± 0.6 events/15 min) and 45-60 min after clemizole exposure (1.3 ± 0.3 events/15 min). Ictal events are noted by open red circles above traces. C, Individual value plot of ictal event frequency at baseline (0-15 min) and 45-60 min after continuous bath application of clemizole (TRZ, n = 14). Independent experiment pairs are indicated by the connected line. \*\*\**p* < 0.0001. D, Representative 60 min LFP traces for *stxbp1b* mutant zebrafish at baseline (8.2 ± 1.4 events/15 min) and 45-60 min after trazodone exposure (1.4 ± 0.4 events/15 min). Ictal events are noted by open red circles above traces. Student’s unpaired *t* tests.

## 4 Discussion

Here we show that repurposed drugs (clemizole and trazodone) can exert powerful antiseizure effects in a larval zebrafish model of STXBP1-related disorders. Our data also confirm a pharmacoresistant phenotype for *stxbp1b* mutant zebrafish, consistent with clinical classifications in this patient population. Although clemizole and trazodone were initially marketed as an antihistamine and antidepressant, respectively, recent data suggests that both exert antiseizure activity via modulation of serotonin signaling (20).

Mutations in STXBP1 were initially identified in patients diagnosed with Ohtahara syndrome e.g., catastrophic early infantile encephalopathy characterized by severe seizures, psychomotor retardation and a high mortality rate (5, 35). Subsequently, mutations were identified in additional neurodevelopmental disorders such as atypical Rett syndrome, Dravet syndrome, West syndrome and Lennox-Gastaut syndrome (8). Overall, STXBP1-related disorders encompass a clinical phenotypic spectrum with nearly all patients exhibiting some form of pharmacoresistant epilepsy. Although patients do not experience seizure freedom with any AED, in single cases, a therapeutic response to vigabatrin, carbamazepine, phenobarbital, valproic acid or levetiracetam have been reported (36–45). In rare examples, ketogenic diet (46) and surgery (38) have also shown some success. However, for most patients, three or more AEDs are used concurrently with little to no seizure control. In the present study, we evaluated prototype AEDs with different putative mechanisms of action (31, 32): (i) PHT and VPA block voltage-gated sodium channels, (ii) LEV mediates synaptic vesicle release, (iii) DZP is a benzodiazepine which enhances GABA_A_-receptor mediated inhibition and (iv) ESX is a low-voltage activated calcium channel blocker commonly used to control absence seizures. Although some *stxbp1* mutant larvae responded to VPA, none of these FDA-approved drugs significantly reduced seizure activity which indicates that this model mirrors human pharmacoresistance seen in this patient population.

Clemizole and trazodone were discovered in a two-stage phenotype-based screen using convulsive behavior as a high-throughput first-pass surrogate for spontaneous seizures followed by LFP monitoring to demonstrate frank suppression of electrographic seizure events. Both repurposed drugs have already advanced to clinical applications (47)(https://clinicaltrials.gov/ct2/show/NCT05066217). This approach utilized a zebrafish voltage-activated sodium channel mutant (*scn1lab*) representing a severe childhood genetic epilepsy condition e.g., Dravet syndrome (DS)(16). Subsequent studies incorporating medicinal chemistry and receptor binding assays identified a potential mechanism of action via binding to a 2B serotonin receptor subunit (20). Fenfluramine, also successfully identified in *scn1lab* mutant zebrafish (17–19) and recently approved by the FDA for treatment of DS, blocks serotonin reuptake. Here, baseline frequency of convulsive seizure events in *stxbp1b* is lower than that observed with the *scn1lab* mutant line precluding high-throughput locomotion-based assays. Behavioral seizure assays are also inherently variable and prone to detection of false-positive drugs that reduce convulsive movements simply by acting as sedative or muscle relaxant (33). Although seizures often have clear behavioral components, a “gold standard” classification of epilepsy in any species is based on detection of electrographic seizure events. Indeed, the literature on drugs purported to have antiseizure activity in larval behavioral assays, but not confirmed using electrophysiology readouts, includes several dozen drugs (48); none of which have advanced to clinical applications. It was also not possible to directly compare our findings with data from *Stxbp1* mutant mice (12, 49) or *STXBP1/Munc18-1* worms (50, 51) as none of the drugs used here have been tested against seizure activity in these models. For these reasons, our investigations focused on suppression (or not) of spontaneous ictal-like events as a sensitive, low-throughput, antiseizure outcome measure. This strategy clearly identified clemizole and trazodone as exerting powerful antiseizure activity in a zebrafish model of STXBP1-related disorders. These findings are consistent with previous demonstrations that modulation of 5HT receptor subtypes can effectively reduce seizures in preclinical zebrafish models. Although some of these drugs also target dopamine, sigma opioid or other 5HT receptor subunits, the most coherent unifying explanation for antiseizure activity reported for clemizole (16), trazodone (20), lisuride (52), fenfluramine (17, 19), and lorcaserin (18, 20) would be an action on the 2B serotonin receptor subunit (53). These observations, previously restricted to *SCN1* are extended here to an entirely new genetic epilepsy population i.e., STXBP1-related disorders. In conclusion, our preclinical results represent an exciting first step in identification of new therapeutic avenues and warrant future investigations, as well as exploratory clinical studies, in these patients.

## Author Contributions

Electrophysiology experiments were performed by M.M. and S.C.B. Data analysis was performed by S.C.B. The design, conceptualization, interpretation of data, data analysis, and writing of the manuscript was done by S.C.B.

